# A Novel Approach to Investigate Subcortical and Cortical Sensitivity to Temporal Structure Simultaneously

**DOI:** 10.1101/2020.03.03.968404

**Authors:** Sonia Y. Varma, David Purcell, Sangamanatha A. Veeranna, Ingrid S. Johnsrude, Björn Herrmann

**Affiliations:** Department of Psychology, Brain and Mind Institute, University of Western Ontario, Canada; School of Communications and Sciences and Disorders, University of Western Ontario, Canada; Rotman Research Institute, Baycrest, Toronto, ON, Canada; Department of Psychology, University of Toronto, Toronto, ON, Canada

**Author notes:** Shared senior authorship (Ingrid S. Johnsrude, Björn Herrmann). Corresponding author: Sonia Varma 1151 Richmond St, London, ON N6A 3K7.

**Keywords:** Brainstem, auditory cortex, adaptation, auditory processing, electroencephalography

## Abstract

Hearing loss is associated with changes at the peripheral, subcortical, and cortical auditory stages. Research often focuses on these stages in isolation, but peripheral damage has cascading effects on central processing, and different stages are interconnected through extensive feedforward and feedback projections. Accordingly, assessment of the entire auditory system is needed to understand auditory pathology. Using a novel stimulus paired with electroencephalography in young, normal-hearing adults, we assess neural function at multiple stages of the auditory pathway simultaneously. We employ click trains that repeatedly accelerate then decelerate (3.5 Hz frequency modulation; FM) introducing varying inter-click-intervals (4 to 40 ms). We measured the amplitude of cortical potentials, and the latencies and amplitudes of Waves III and V of the auditory brainstem response (ABR), to clicks as a function of preceding inter-click-interval. This allowed us to assess cortical processing of frequency modulation, as well as adaptation and neural recovery time in subcortical structures (probably cochlear nuclei and inferior colliculi). Subcortical adaptation to inter-click intervals was reflected in longer latencies. Cortical responses to the 3.5 Hz FM included phase-locking, probably originating from auditory cortex, and sustained activity likely originating from higher-level cortices. We did not observe any correlations between subcortical and cortical responses. By recording neural responses from different stages of the auditory system simultaneously, we can study functional relationships among levels of the auditory system, which may provide a new and helpful window on hearing and hearing impairment.

## 1. Introduction

The multiple stages of the mammalian auditory system are highly interconnected. Feedforward and feedback connections span from the cochlea to cortex (Webster, 1992; Malmierca and Hackett, 2010; Kelly and Wong, 1981; Huffman and Henson, 1990). Changes linked to aging and/or noise exposure have been documented at all stages of the auditory pathway, including the hair cells (Chen and Fechter, 2003), auditory nerve (Kujawa and Liberman, 2009), cochlear nucleus (Caspary et al., 2005), inferior colliculus (Mehraei et al., 2016), and cortex (Mendelson and Ricketts, 2001; Herrmann et al., 2016). Moreover, functional changes at lower levels of the auditory system might elicit changes in higher-level regions (Hickox and Liberman, 2013). For example, damage to the auditory periphery has been associated with a loss of inhibition in cortex and hyper-responsiveness to sound (Salvi et al., 2017; Caspary et al., 2008). Understanding the functional relationships among different stages of the auditory system may assist in characterizing hearing function and dysfunction.

Critically, changes at different stages of the auditory system have been associated with changes in sound and speech perception. For example, subcortical changes in temporal processing have been associated with poor speech perception in noise (Bidelman et al., 2014; Presacco et al., 2016; Ruggles et al., 2012; Ruggles et al., 2011; Mehraei et al., 2016). A loss of inhibition and an increase in the responsiveness of auditory cortex may explain why some older individuals experience sounds at moderate intensities as too loud (Epstein, and Marozeau, 2010; Tyler et al., 2014; Anderson et al., 2012; Knipper et al., 2013; Auerback et al., 2014; Herrmann et al., 2018). Indeed, older individuals manifest larger responses to the regular temporal structure of amplitude- or frequency-modulated sounds (Herrmann et al., 2019; Goossens et al., 2016, Presacco et al 2016). Evaluating multiple levels of the auditory system at once may reveal relations among subcortical and cortical response changes that will illuminate functional dependencies among levels of processing.

Changes in adaptation (reduction in neuronal responsiveness due to repetitive stimulation) occur with age and/or noise exposure (Miller et al., 1994; Li et al., 1994), and may be associated with perceptual deficits (Strait et al., 2011). The interonset interval (IOI) between successive click stimuli determines the time neurons have before another sound is presented, potentially limiting the neural response to that second sound if adaptation is still present. We evaluate adaptation at subcortical levels of the auditory system by measuring how the parameters of the auditory brainstem response (ABR; Jewett and Williston, 1971) change as the interval between successive auditory clicks (IOI) is systematically altered. ABRs are typically evaluated using electroencephalography (EEG). However, electrical potentials recorded from the cochlea (electrocochleography; ECochG) are more sensitive to responses in the auditory nerve and hair cells (Ferraro, 2000) and so we use an integrated EEG-ECochG system to record ECochG simultaneously with EEG.

Neural synchronization to sound (entrained oscillatory activity to low-frequency temporal regularity) is thought to temporally organize cortical excitability, thereby shaping perception (Lakatos et al., 2008; Henry and Herrmann, 2014). Neural synchronization may also support attention to sound over time, and enable predictions about future sounds (Nozaradan et al., 2011; Costa-Faidella et al., 2017; Schroeder and Lakatos, 2009; Nobre and van Ede, 2018; Henry and Herrmann, 2014). It may therefore play an important role in speech comprehension (Peelle and Davis, 2012; Doelling and Poeppel, 2015; Giraud and Poeppel, 2012). Neural synchronization is frequently investigated using frequency- or amplitude-modulated sounds (FM or AM; Henry and Obleser, 2012; Herrmann et al., 2013, 2018; Goossens et al., 2016; Maiste and Picton, 1989; Picton et al., 2003) or speech (Keitel et al., 2017; Ding et al., 2014; Di Liberto et al., 2015), and is thought to reflect activity in auditory cortex (Ding et al., 2014; Giraud and Poeppel, 2012).

Another neural signature of temporal-regularity processing is a low-frequency sustained response (Barascud et al., 2016; Sohoglu and Chait, 2016; Southwell et al., 2017; Gutschalk 2002; Pantev 1996). The sustained response is thought to reflect the detection of regularity in a sound sequence (Barascud et al., 2016; Sohoglu and Chait, 2016; Southwell et al., 2017). The magnitude of the sustained response also depends on whether individuals attend to, or ignore, sound, unlike neural synchronization (Herrmann and Johnsrude, 2018). The sustained response is thought to originate from extra-auditory brain regions including frontal cortex, parietal cortex, and hippocampus (Barascud et al., 2016; Sohoglu and Chait, 2016; Teki et al., 2016), in addition to auditory cortex (Barascud et al., 2016; Pantev et al., 1996; Sohoglu and Chait, 2016; Gutschalk et al., 2002).

Here, we develop a novel stimulus that enables the simultaneous assessment of subcortical and cortical sensitivity to changes in temporal structure. Using an integrated EEG-ECochG system, we evaluate subcortical adaptation by measuring how the parameters of the ABR change as IOI varies while, at the same time, measuring neural synchronization and sustained activity from cortical areas.

## 2. Methods

### 2.1 Participants

Twenty-nine healthy, English speaking adults (mean: 20.41 years, range: 17-28 years, females: 21) participated in the current study. Four additional participants took part in the experiment but were excluded due to >30% of data being contaminated by artifacts (N=2), or having a Wave V latency that was more than two standard deviations longer the mean across all subjects (N=2). Participants reported no neurological diseases or hearing problems. They gave written consent prior to the experiment and were either paid $5 CAD per half hour for their participation or were given course credits for their introductory psychology course. All participants had an average (across both ears) pure-tone threshold of 25 dB HL or less, assessed using pure-tone audiometry at octave frequencies between 0.25-4 kHz. This experiment was conducted in accordance with the Declaration of Helsinki, the Canadian Tri-Council Policy Statement on Ethical Conduct for Research Involving Humans (TCPS2-2014) and was approved by the non-medical research ethics board (NMREB) of the University of Western Ontario (protocol ID: 106570).

### 2.2 Acoustic stimulation and procedure

Prior to the EEG recording, otoscopy was performed on each participant to confirm that the tympanic membrane was intact and to ensure that the ear canal was not occluded. All sounds in the current experiment were presented to the right-ear via an Etymotic ER1 earphone, using a FIREFACE 400 sound card controlled by Psychtoolbox (version 3) in MATLAB (MathWorks Inc.). For each participant, the sensation level for click trains was determined using a method of limits procedure estimating the 50% hearing threshold (for a detailed description see Herrmann and Johnsrude 2018a, b), and all experimental stimuli were presented at 60 dB above the individual hearing threshold.

Prior to the main experiment, EEG was recorded while participants listened to a validation stimulus similar to that used in standard clinical protocols. This stimulus consisted of 4000 clicks (0.1 ms duration, rectangle function) that were presented with an inter-onset-interval (IOI) of 0.088 s (11.3 Hz). Clicks were presented with alternating polarity at every 10^th^ click (10 clicks of one polarity followed by 10 clicks of the opposite polarity etc.) in order to minimize stimulus artifacts and the cochlear microphonic in the trial average (Eggermont, 2017). The validation stimulus lasted about 6 min. These recordings were used to confirm the sensitivity of our recording system to peripheral and subcortical potentials (Wave I/CAP, Wave III, and Wave V).

In the main study, two stimulus conditions were presented: an unmodulated control stimulus and a frequency-modulated stimulus (Figure 1). The unmodulated control stimulus (Figure 1a) consisted of 140 clicks (0.1 ms duration, rectangle function, alternating polarity at every 10 clicks) presented at a fixed IOI of 0.04 s, resulting in a ~5.64 s stimulus. The frequency-modulated stimulus (Figure 1b/c) was a click train in which the IOIs changed from 0.04 to 0.004 s and back again, creating a modulated click pattern (18 IOIs; logarithmically spaced; see also Herrmann et al., 2016). The continuously accelerating and decelerating IOI resulted in a 3.5 Hz frequency-modulated sound, covering frequencies from 25 Hz (0.04 s IOI) to 250 Hz (0.004 s IOI). The modulated part of the stimulus consisted of 16 accelerating-decelerating cycles and was preceded by 1.08 seconds without modulation, so identical to the unmodulated control stimulus (27 clicks with an IOI of 0.04 s). The identical onsets for both stimuli allowed us to observe how responses diverge between the two conditions once the modulation began. The total duration of the stimulus incorporating frequency modulation was also ~5.64 seconds. Stimuli were presented in alternating polarity.

**Figure 1:**
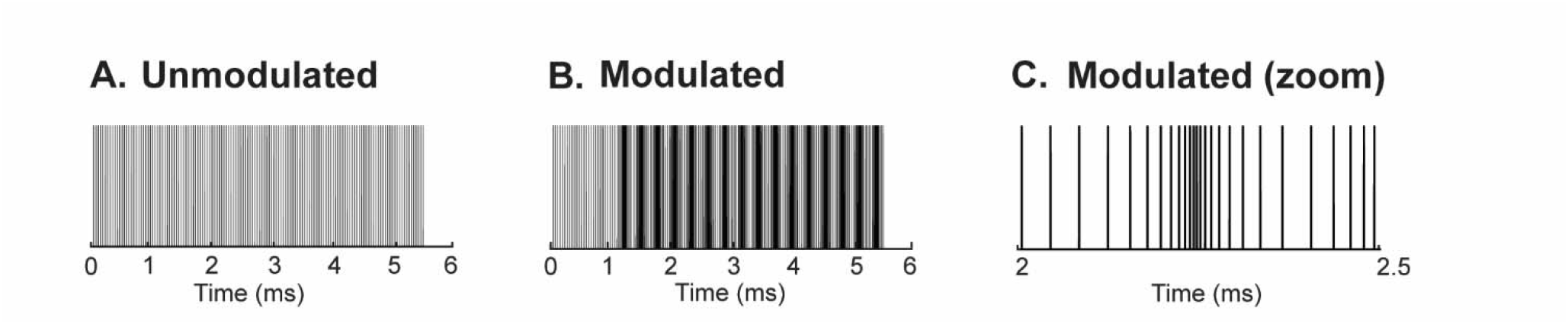
Acoustic stimuli of the main experiment. **A:** Clicks over time for the unmodulated stimulus. Each vertical line represents one click. **B:** Clicks over time for the 3.5 Hz frequency-modulated stimulus. The frequency modulation starts at 1.08 s. **C:** Zoomed in representation of the frequency-modulated stimulus shown in panel B. Each vertical line represents one click.

In each of six blocks, the unmodulated and modulated stimulus were each presented 24 times, resulting in 144 trials for each condition. Stimuli were presented in pseudo-random order such that the same stimulus could be presented a maximum of three times in direct succession. Sounds were separated by a silent inter-stimulus interval of ~1.96 seconds.

During sound stimulation in the validation paradigm and experimental blocks, participants watched a muted movie of their choosing with subtitles. Participants were instructed to ignore the acoustic stimulation and watch the movie.

### 2.3 EEG recording

Participants were seated in comfortable chair in a single-walled sound-attenuating booth (Eckel). Electroencephalograms were recorded using a 16-channel ActiveTwo BioSemi system (Cortech Solutions, Inc.). The sampling rate was 16,384 Hz (a 3334 Hz online low-pass filter was used to avoid aliasing; this filter is built into the BioSemi system). Additional surface electrodes were placed on both mastoids and earlobes. Channels were referenced online to the CMS electrode located adjacent to CZ. The EEG recording setup additionally included simultaneous extra-tympanic ECochG recording capabilities. ECochG was implemented using an individual active electrode custom-made from a BioSemi electrode that was connected to a clinical ECochG gold-foil tiptrode (Etymotic Research Inc.). This tiptrode serves as an electrode but is also a ER1-compatible eartip, delivering the stimulus to the ear via silicone tubing (52 cm total tube length) connected to the ER1 system. The gold-foil tiptrode was inserted into the participant’s right ear canal and ECochG signal referenced online to the CMS electrode.

### 2.4 EEG analysis

Offline data analysis was carried out using MATLAB software (MathWorks, Inc.). Raw data were notch filtered with a 60-Hz band-stop, elliptic filter to suppress line noise.

#### Analysis of subcortical responses

Data from 16 scalp electrodes were re-referenced to the averaged mastoids, and the data from the in-ear tiptrode was re-referenced to the contralateral (left) ear lobe. Raw data were high-pass filtered at 130 Hz (2743 points, Hann window) and low-pass filtered at 2500 Hz (101 points, Hann window). Data were divided into epochs ranging from −2 ms to 10 ms time-locked to the click onset at 0 ms. Epochs were rejected if the amplitude range exceeded 60 µV in the Cz or C3 electrodes. These electrodes were selected because sounds were presented monaurally to the right ear only, eliciting the most pronounced responses around the center to left side of the scalp (Woods et al., 1985). The rejection criterion was applied only to the time-window following stimulus onset (0-10 ms), so as to not reject trials based on any artifact which may precede the click onset, since this would be canceled in the across-trial average due to the alternating polarity of the stimuli.

To explore whether our stimulus elicited similar subcortical responses compared to the validation stimulus (with clicks presented at a slower rate), responses to individual clicks in the unmodulated control stimulus (from the experimental blocks; IOI of 0.04 s) were averaged and compared to averaged responses to clicks in the validation stimulus (IOI of 0.088 s).

To examine the sensitivity of subcortical responses to IOI (i.e., to measure adaptation), neural responses elicited by individual clicks in the modulated section of the frequency-modulated sound were extracted and sorted according the IOI directly preceding each click. Single-trial responses for a unique IOI (irrespective of whether it was presented in accelerating or decelerating sections) and its immediately adjacent neighbors on either side (i.e., shorter and longer intervals) were binned and averaged (Ingham and McAlpine, 2005; Herrmann et al., 2016). Adjacent intervals were included to increase power.

We investigated Auditory Brainstem Responses (ABRs) using the averaged signal across Cz and C3 electrodes (for Wave V), and the signal from the in-ear tiptrode (for Waves I and III). We were interested in Wave I, Wave III, and Wave V, since these responses are thought to originate from different subcortical generators (auditory nerve fibers, cochlear nuclei, and inferior colliculi, respectively; Picton, 2010). Waves were identified based on baseline-to-peak latency and morphology (Picton, 2010). However, very short-latency responses (<3 ms), characteristic of Wave I, were not observed in the ABR or in-ear tiptrode and were thus not included in further analyses. The reason for the absence of Wave I is unclear but could in part be related to the relatively low sound intensity of 60 dB SL (equal to about 80 dB SPL; Moore et al., 2004) utilized in the current study compared to previous work (Santarelli et al.,2002; McMahon et al., 2002). Sounds in the current study could not be presented at a higher intensity since the fast click rates we used would have been perceived as uncomfortably loud. Waves III and V were present in the majority of participants and the characteristics of these components are analysed.

The Wave III amplitude and latency were extracted from the in-ear tiptrode (ECochG) recordings, separately for each participant. The tiptrode was used in order to maximize the size and reliability of the responses. Only participants who displayed a Wave III to the validation stimulus (N=21) were included in this analysis. The Wave V amplitude and latency were extracted from the averaged Cz and C3 electrodes (all N=29 participants were included). For each participant, a linear function was separately fit to amplitude and latency data as a function of IOI (independently for Wave III and Wave V). Positive slopes would indicate an increase in amplitude/latency with increasing IOI and vice versa. The slope of the linear fit, for amplitude and latency data for Waves III and V, were tested against zero using one-sample t-tests.

#### Analysis of cortical responses

We evaluated neural synchronization of cortical responses to the frequency-modulated stimuli, using the unmodulated stimuli as a control. We evaluated both inter-trial phase coherence (ITPC; Lachaux et al., 1999) and the sustained response as markers for synchronization.

Data from the 16 scalp electrodes were re-referenced to the averaged signal from the two mastoid electrodes. Data were low-pass filtered at 22 Hz (2001 points, Kaiser Window), down-sampled from 16,384 Hz to 1024 Hz, and high-pass filtered at 0.7 Hz (2449 points, Hann window). Data were divided into epochs ranging from −1 s to 6.6 s time-locked to stimulus onset at 0 s (first click of a stimulus train). Independent Components Analysis (ICA; runica method, Makeig at al., 1996; logistic informax algorithm, Bell and Sejnowski, 1995) was computed using Fieldtrip software (v2017b). Components containing eye blinks were rejected, and the data were then projected onto the original electrodes. Additionally, trials were rejected if the amplitude range exceeded 200 µV in any of the 16 electrodes.

Neural synchronization was investigated by calculating ITPC. A fast Fourier transform (including Hann window taper and zero padding) was calculated for each trial and channel during the time window in which regularity (frequency modulation) could occur (1.08 to 5.64 s). The resulting complex numbers were divided by their respective magnitudes to obtain unit amplitude. The normalized complex numbers were averaged across trials, and their absolute value was used as ITPC. ITPC takes on values between 0 and 1, with 0 reflecting no coherence and 1 reflecting maximum coherence. ITPC was calculated for frequencies within the range of 0 to 12 Hz and averaged across the fronto-parietal electrode cluster (Fz, F3, F4, Cz, C3, C4, Pz, P3, P4). In order to compare neural synchronization between conditions (unmodulated vs frequency-modulated sounds) and thus determine whether the neural activity synchronized with the frequency modulation in sounds, the ITPC magnitude was extracted around the 3.5 Hz (3.45 to 3.55 Hz) stimulus frequency modulation rate. ITPC was compared between conditions using a dependent samples t-test.

The analysis of the sustained response involved averaging single-trial time courses separately for each condition (unmodulated and frequency-modulated). High-pass filtering was omitted for this analysis, since the sustained response is a low-frequency signal (Barascud et al., 2016; Southwell et al., 2017, Herrmann and Johnsrude, 2018). Rejection of components with blinks was based on the independent components analysis using the high-pass filtered data. Responses were baseline-corrected by subtracting the mean amplitude of the pre-stimulus time window (–1 s to 0 s) from the amplitude at each time point. Signals were averaged across a fronto-parietal electrode cluster (Fz, F3, F4, Cz, C3, C4, Pz, P3, P4; see Herrmann and Johnsrude 2018), and the mean amplitude was calculated within the time window in which the frequency modulation was present in the modulated condition (1.08 to 5.64 s). The amplitude of the sustained response was compared between conditions (unmodulated vs frequency-modulated sounds) using a dependent samples t-test.

### 2.5 Correlations

In order to investigate the relation between measures of neural activity (thought to originate from different neural generators), Pearson’s correlations were calculated among the following metrics: Overall (averaged across all IOIs) Wave III and V latency and amplitude, slopes that relate Wave III and V latency and amplitude to IOIs, neural synchronization (i.e., ITPC difference between modulated and unmodulated stimuli), and the magnitude of sustained response (i.e., amplitude difference between modulated and unmodulated stimuli). The false discovery rate correction (Benjamin and Hochberg, 1995) was used to control false-positive rate.

## 3. Results

### 3.1 Subcortical responses are similar between experimental and validation protocol

In order to confirm that our stimulus paradigm is sensitive to neural responses that are commonly obtained in clinical settings, we compared responses to clicks in the unmodulated stimulus (presented in the experimental blocks) to responses to clicks in the validation protocol. Figure 2, depicting the grand average time courses, shows that both stimulus protocols elicit reliable Waves III and V peaks at approximately 4 ms and 6 ms, respectively. The latencies of Waves III and V were slightly delayed in response to the experimental stimulus, compared to the validation stimulus (t_20_= - 2.78, p= 0.011, and t_28_= −2.75, p= 0.012, respectively). This was expected as the clicks in the experimental stimulus are presented at a much faster rate (11.3 Hz vs 25 Hz; Hood, 1998; Burkhard and Sims, 2001; Harkins, McEvoy, and Scott, 1979). The amplitudes of Waves III and V did not differ between the two stimulus types (t_20_= 1.89, p= 0.078, and t_28_= 0.87, p= 0.093, respectively). These results demonstrate that our experimental protocol and the ECochG electrode are sensitive to subcortical responses.

**Figure 2:**
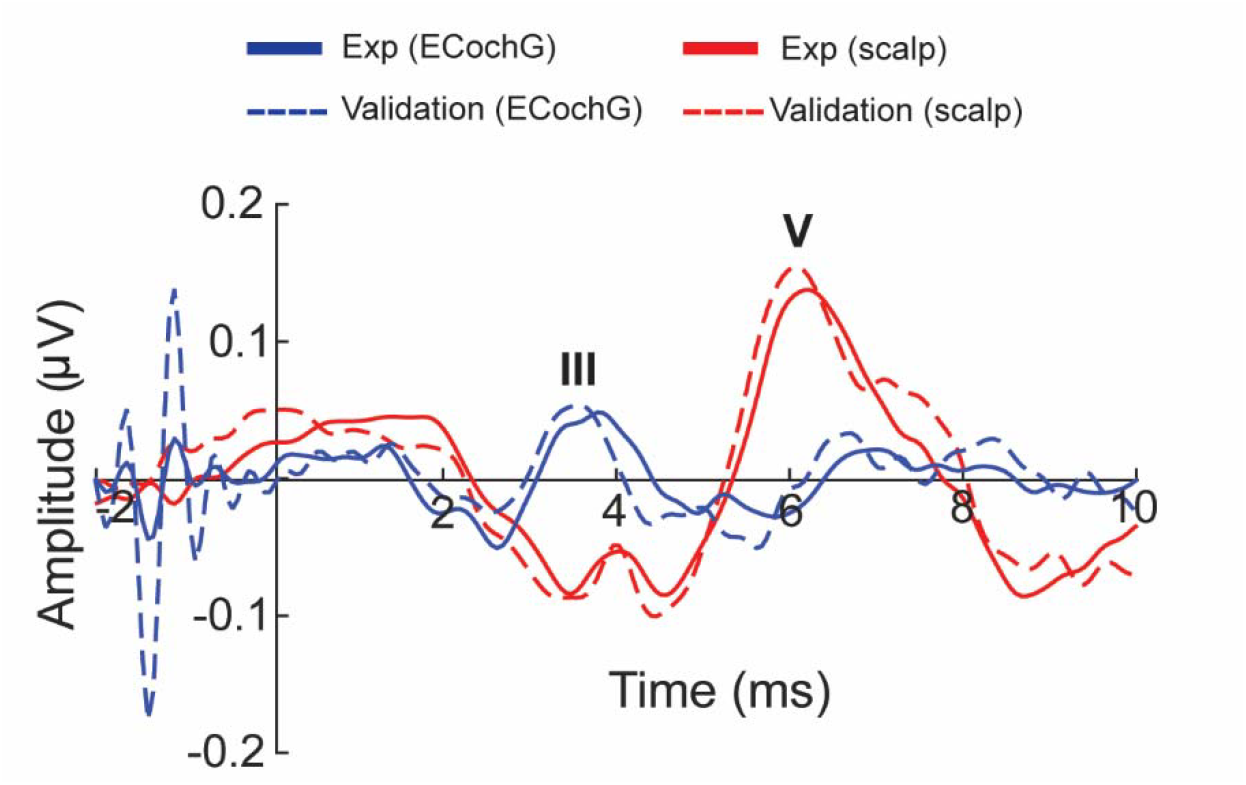
Responses to clicks for isochronous IOIs in the validation and experimental (unmodulated) control stimuli. Red lines show time courses from the averaged signals across C3 and Cz electrodes, whereas blue lines reflect the time courses for the in-ear tiptrode (ECochG; solid for experimental protocol, IOI = 0.04 s; dashed for validation protocol, IOI = 0.088 s).

### 3.2 Latencies, but not amplitudes of Wave III and V are sensitive to IOI

Figure 3A shows the ECochG time courses elicited by the clicks, separately for different IOIs. Individual Wave III amplitudes and latencies were extracted for each participant, and linear functions were fitted to the data relating amplitudes and latencies to IOI. The pattern of results is broadly consistent with adaptation: Wave III amplitude tended to increase in each subject with increasing IOI (slopes of the linear fit approached a significant difference from zero; t_20_ = 1.9489, p = 0.0655; Figure 3B). Furthermore, latencies decreased as the IOI increased: the slopes of the linear function fit was significantly smaller than zero (t_20_ = −2.4248, p = 0.0249; Figure 3C). Since the Wave III latency peaks at ~4 ms, which is equal to one of the IOIs used, and may lead to overlapping responses, the same analysis was performed using IOIs greater than 7 ms only. This cut-off eliminated IOIs that were shorter than the latest peak in the ECochG waveform. In this analysis, Wave III amplitude increased reliably with increasing IOI (slopes greater than zero: t_20_ = 2.3057, p = 0.0320), and latencies reliably decreased with increasing IOI (slopes smaller than zero: t_20_ = −2.217, p = 0.0384). Amplitude and latency of Wave III are sensitive to the interval between clicks.

**Figure 3:**
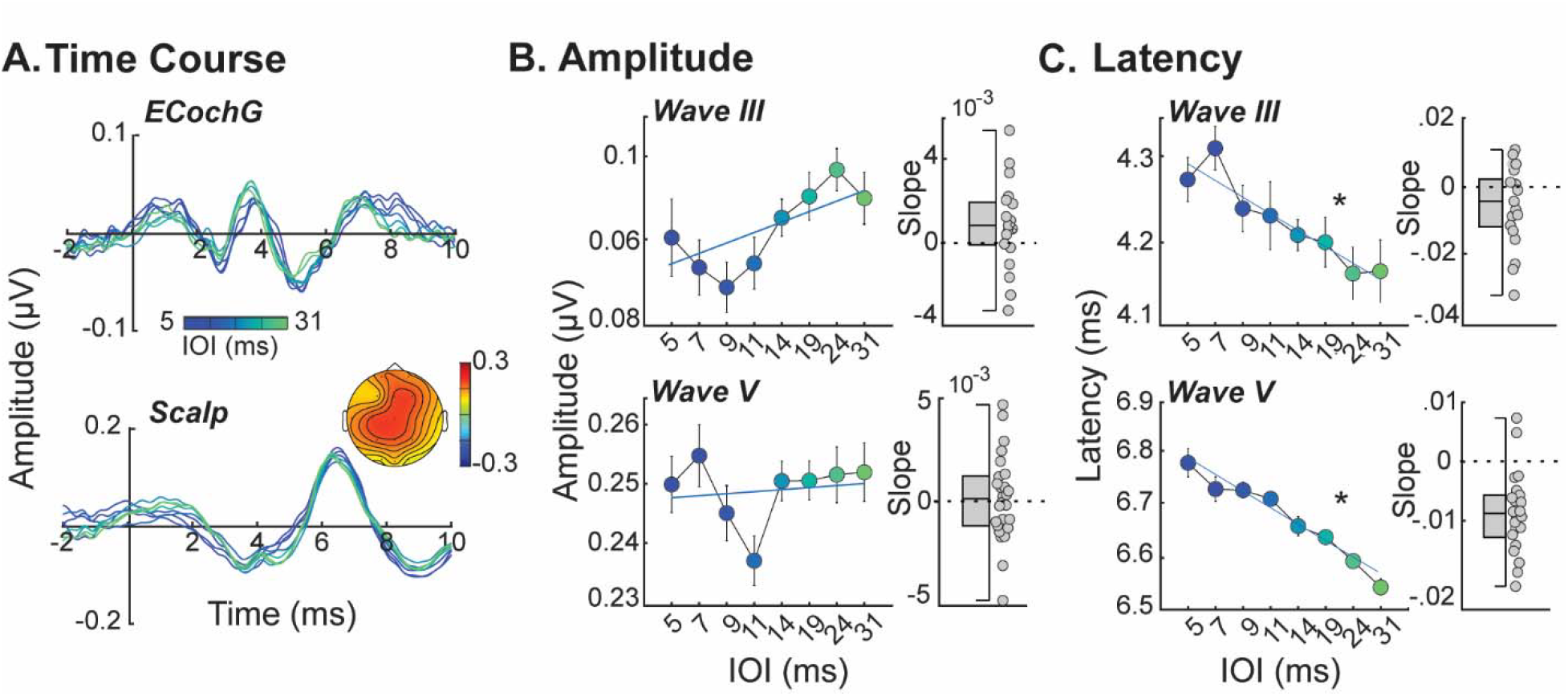
Effects of inter-onset-interval (IOI) on subcortical responses. **A:** Average time-courses of Wave III (N=21) from the ECochG tip-trode (top) and Wave V (N=29) from the averaged Cz and C3 scalp electrodes for each of 8 unique inter-onset intervals (IOIs). Color bar corresponds to the IOI preceding the clicks used to produce each time-course. Topography in lower panel reflects a time-window of ± 1 ms around the peak Wave V latency for all subjects and IOIs. **B:** Wave III (top panel) and Wave V (bottom panel) amplitudes as a function of IOI, averaged over subjects. Individual slope values (taken from each participant’s linear fit) on the right-hand side. **C:** As Panel B, but for Wave III and Wave V latencies. *p ≤ 0.05 indicates a significant difference from zero.

Figure 3A (bottom panel) shows the response time courses for the average across Cz and C3, and a Wave V peak around 6.5 ms. The mean slope for the linear function relating Wave V amplitude to IOI did not differ from zero (t_28_ = 0.4232, p = 0.6754; Figure 3B). In contrast, Wave V latencies reliably decreased with increasing IOI (slopes smaller than zero: t_28_ = −7.519, p < .001; Figure 3C). The Wave V latency peaked at ~6.5 ms, which is longer than some IOIs. To eliminate potential influences from overlapping responses, the same analysis was performed using IOIs greater than 9 ms only. This cut-off eliminated IOIs that were shorter than the latest peak in the ABR waveform. The exclusion of the IOIs below 9 ms did not alter the findings. The mean slope of Wave V amplitude was not significantly different from zero (t_28_ = 1.5484, p = 0.132), while the mean slope of Wave V latency was significantly smaller than zero (t_28_ = −7.951, p < .001). These results show that Wave V latency is sensitive to IOI.

### 3.3 Neural activity synchronizes with frequency modulation in sounds

Figure 4A shows ITPC for the unmodulated control stimulus and the 3.5-Hz frequency-modulated stimulus. ITPC at 3.5 Hz was significantly larger for the frequency-modulated compared to the unmodulated stimulus (t_28_ = −7.58, p < .001), indicating synchronization of neural activity with the imposed frequency modulation. Due to the large difference in variability between the modulated and unmodulated conditions, a non-parametric Wilcoxon Signed-Ranks test was also performed, which confirmed that ITPC at 3.5 Hz was significantly larger for the frequency-modulated compared to the unmodulated stimulus (Z_28_ = −4.70, p < .001).

**Figure 4:**
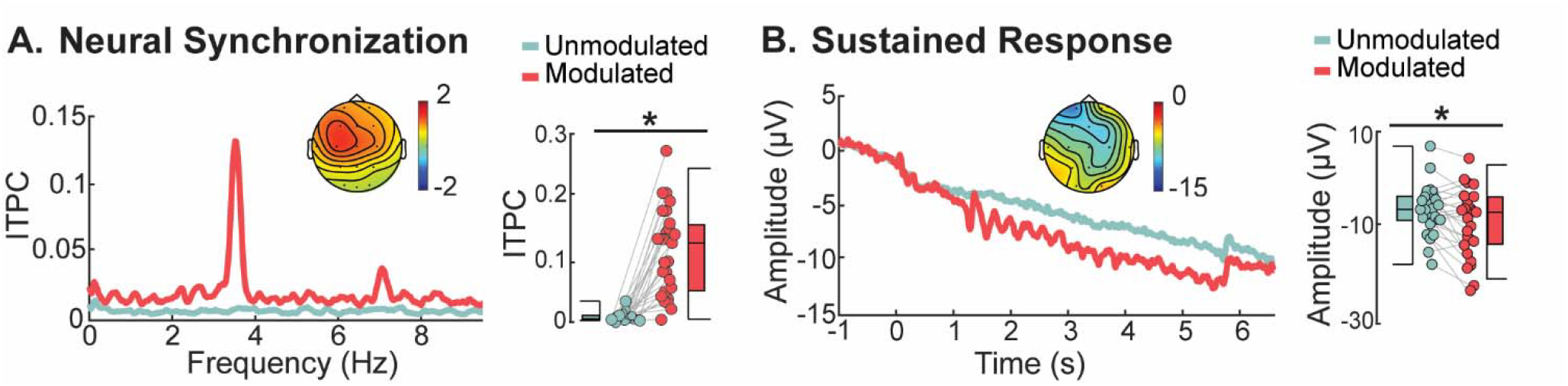
Cortical responses to the 3.5-Hz frequency-modulated stimulus and the unmodulated control stimulus. **A:** Inter-trial phase coherence (ITPC) spectra are displayed on the right. IPTC was averaged across the 3.45-3.55 Hz frequency window in order to evaluate neural synchronization at the stimulus modulation frequency (3.5 Hz). IPTC values for individual participants are depicted on the right for both conditions (lines connect points from the same individual). *p ≤ 0.05. Topography reflects ITPC in the frequency window of 3.45 to 3.55 Hz. **B:** Sustained activity for unmodulated and frequency-modulated stimulus conditions. Amplitudes were extracted from the time window during which the frequency modulation was present in the modulated stimulus (1.08 s to 5.64 s), and individual data are displayed on the right (lines connect points from the same individual. *p ≤ 0.05. Topography shows mean amplitude in the time window of 1.08 s to 5.64 s.

### 3.4 Sustained activity in the cortex is enhanced by temporal regularity

Sustained activity in the time window of 1.08 to 5.64 s (during which the frequency modulation was present in the modulated stimulus) was larger (i.e., more negative) for the frequency-modulated compared to the unmodulated stimulus (t_28_ = 2.725, p = 0.013). A non-parametric Wilcoxon Signed-Ranks test was also performed, which confirmed that sustained activity in the time window of 1.08 to 5.64 s was larger (i.e., more negative) for the frequency-modulated compared to the unmodulated stimulus (Z_28_ = 2.368, p = 0.018).

### 3.5 Correlations among neural measures

The previous sections demonstrate that subcortical responses are sensitive to IOI (Wave III and V), and cortical responses are sensitive to the presence of frequency modulation (neural synchronization and sustained activity). Wave III and V are thought to originate from the cochlear nucleus and inferior colliculus, respectively (Picton, 2010), whereas neural synchronization to low-frequency modulations (i.e. 3.5 Hz) is thought to originate from auditory cortex (Herrmann et al. 2013; Keitel et al., 2017) and regularity-related sustained activity is thought to be generated by auditory and higher-level cortices (Barascud et al., 2016; Gutschalk et al., 2002; Pantev et al., 1996; Teki et al., 2016). We calculated correlations among all our neural measures to explore potential relations among different stages of the auditory system in sensitivity to temporal structure in sound.

Figure 5A shows the correlation matrix (Pearson’s correlations, r) for our neural measures: mean amplitudes and latencies for Wave III and Wave V, slopes of the linear function relating Wave III and Wave V amplitudes and latencies to IOI, ITPC difference between the modulated and unmodulated conditions, and sustained response difference between the modulated and unmodulated conditions.

**Figure 5.**
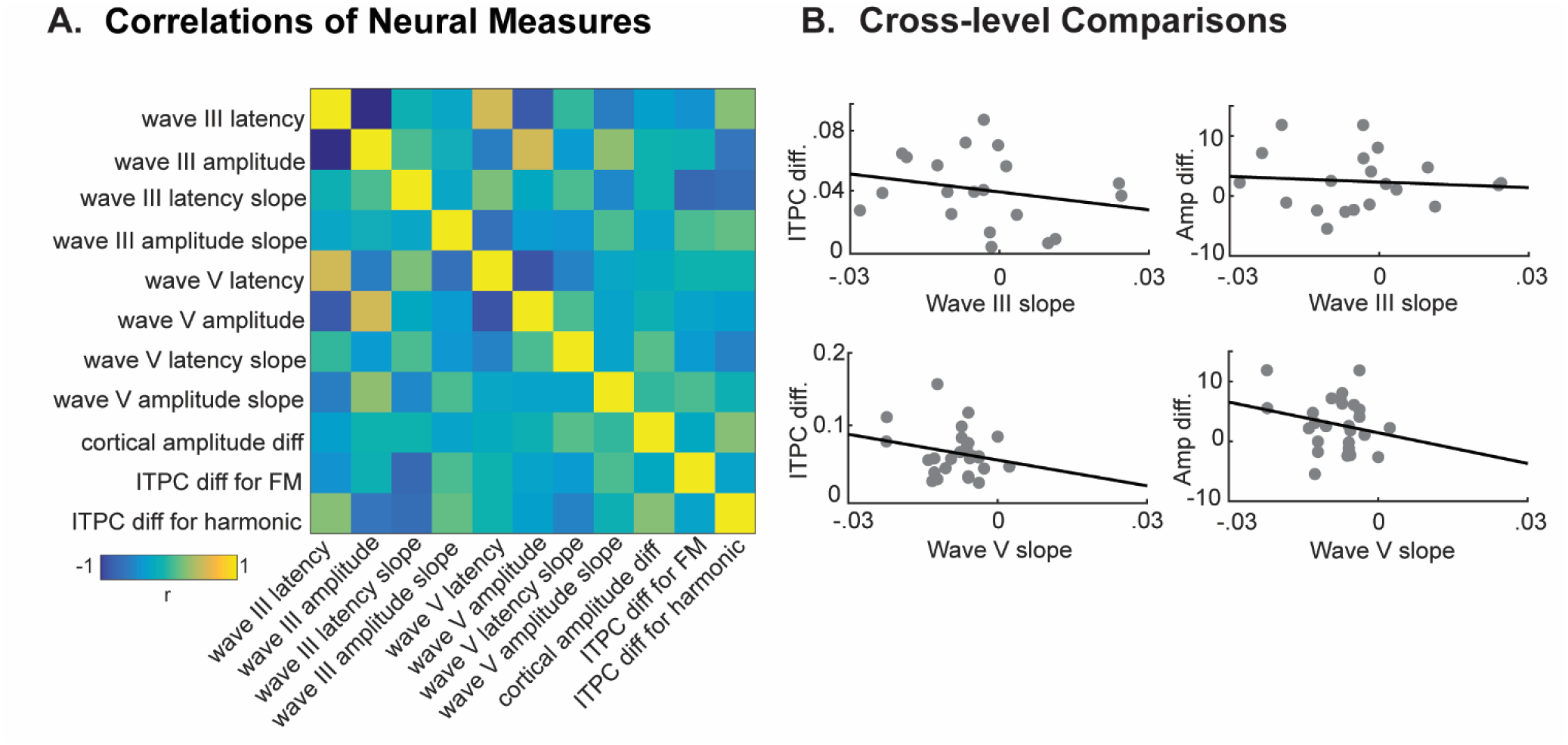
Correlations among neural measures. **A**. Full correlation matrix. **B**. Sample of cross-level correlations between: LEFT: slope of subcortical (Wave III, top; Wave V, bottom) latency change as a function of IOI and RIGHT: cortical processing of frequency modulation (sustained response amplitude difference between conditions, top; ITPC difference at 3.5 Hz between conditions, bottom).

No across-level (subcortical vs cortical) correlations were significant (all p_FDR_ > 0.05). The only significant correlations were between the latency and amplitude of Wave III (r = −0.77, p_FDR_ = 0.0015), and of Wave V (r = −0.64, p_FDR_ = 0.0044), suggesting that individuals with prolonged wave latencies also display reduced wave amplitudes.

## 4. Discussion

We explored a novel stimulus protocol in which clicks were presented with steadily decreasing and then increasing IOIs such that the stream was frequency modulated at 3.5 Hz. This allowed us to simultaneously and noninvasively assess the sensitivity of different stages of the human auditory system to changes in temporal structure. We investigated the relationship between temporal processing in subcortical (cochlear nucleus and inferior colliculus) and cortical auditory structures. Subcortically, the latency of Wave III and Wave V were sensitive to IOI, with amplitude of Wave III also exhibiting sensitivity. Cortical sensitivity to the 3.5-Hz frequency modulation in the same stimulus was demonstrated by neural synchronization and sustained neural activity. No simple correlations were found between subcortical and cortical responses, indicating that any relationship between levels may be mediated by additional factors, or is complex, and potentially non-linear. The current study provides a novel method that enables the simultaneously assessment of subcortical and cortical sensitivity to temporal structure in sounds.

### 4.1 Comparison of novel stimulus with a standard clinical protocol

In the current study, our new stimulus protocol elicits Waves III and V of the ABR as reliably as does a standard protocol (our validation stimulus). These results are consistent with a large literature on Wave III and V, reporting reliable peaks at approximately 4 ms and 6 ms, respectively (Hall, 2007; Picton, 2010). This confirmed that our unmodulated stimulus (0.04 s IOI) elicits reliable subcortical responses despite faster stimulation rates compared to a standard clinical stimulus (0.088 s IOI).

### 4.2 Absence of Wave I

Neither the validation nor experimental stimulus elicited a reliable Wave I response, which originates from auditory nerve fibers (Moore, 1987). Even the ECochG electrode, which was placed closest to the generator in the auditory nerve (i.e., in the ear canal), appeared to be insensitive to auditory nerve fiber responses. The magnitude of human Wave I (or the ECoG compound action potential; CAP) is typically small (Sohmer and Feinmesser, 1973), which can make it difficult to record using extra-tympanic ECochG. The relatively low click intensity (60 dB sensation level) may have contributed the absence of Wave I. In clinical setups, sound levels are often around 90 dB peak-to-peak equivalent SPL (Ferraro, 2000), whereas our 60-dB sensation level probably corresponds to ~80 dB peak-to-peak equivalent SPL (Moore et al., 2004). We were, however, unable to increase the sound intensity further as the fast click presentation rate would have made the sound presentation uncomfortably loud.

### 4.3 Subcortical Sensitivity to Changes in Temporal Structure

Subcortical sensitivity to IOI was evaluated by measuring latency and amplitude of Waves III and V as a function of IOI. Wave III and V latencies decreased with increasing IOIs, whereas amplitudes were less affected by IOI. This is consistent with many previous observations, showing that Wave III and V latencies are sensitive to IOI (Hood, 1998; Burkhard and Sims, 2001; Harkins et al., 1979), whereas Wave III and V amplitudes are generally less sensitive to IOI (Hood, 1998; Lasky, 1984; Suzuki, Kobayashi and Takagi, 1986). Previous research employed a range of IOIs from 0.1 s to 0.01 s (10 Hz to 100 Hz; Lasky, 1984; Suzuki et al., 1986). In contrast, the current investigation used much shorter IOIs, ranging from 0.04 s to 0.004 s (25 Hz to 250 Hz). When we excluded the very shortest IOIs, our results did not change appreciably, except that sensitivity of Wave III amplitude to IOI was observed only after exclusion of very short IOIs (which might allow contamination from overlapping responses). Wave V amplitude was not sensitive to IOI, even when the shortest IOIs were excluded: this is consistent with several previous studies documenting that Wave V amplitudes are less affected by stimulus rate compared to those of other components (i.e. Waves 1 and III) Jewett and Williston 1971; Picton et al. 1974, 1981; Terkildsen et al. 1975; Thornton and Coleman 1975; Hyde et al. 1976; Pratt and Sohmer 1976; Chiappa et al. 1979; Van Olphen et al. 1979).

A likely neurophysiological mechanism underlying sensitivity to IOI is neural adaptation. Adaptation refers to a reduction in neuronal responsiveness due to repetitive stimulation (Li et al.,1993; Miller and Desimone, 1994). With increasing IOIs, neurons can recover for longer before the next sound stimulation, thereby decreasing the response latency (Wave III and V) or increasing amplitude (Wave III) to sound. This interpretation is consistent with single-neuron recordings from the inferior colliculus in rats, which demonstrate both reduced firing rates and longer first-spike latencies to repetitive stimulation (tone bursts with 0.3 s IOI; Herrmann et al., 2015). These changes may occur because sounds are presented while neurons sensitive to the sound are in a refractory state, resulting in a reduced firing rate across a population and reduced magnitude of EEG signal. Peak latency would be shorter when IOIs are long, because most of the population can respond immediately resulting in a coherent and measurable response. In contrast, when IOIs are shorter, a coherent response is not possible until a sizable proportion of cells have recovered.

It is not clear why adaptation is not evident in the amplitude of Wave V, since amplitudes should reflect firing rate more directly. While our findings are consistent with some previous reports of IOI affecting Wave V latency, but not amplitude (Lasky, 1984; Suzuki et al., 1986), a considerable number of investigations have found Wave V amplitude to be affected by presentation rates over 40 Hz (thought still less than Wave V latency; Picton et al., 1981). The discrepancy between our findings and previous ones (that find IOI effects on amplitude) may be because the individual IOIs were only presented in the context of a relatively slow frequency modulated rate (3.5 Hz). This may have provided enough time at slower IOIs for some brainstem neurons to recover and, thus, limit effects on amplitude, but not latency. It remains unclear why ABR latency is more sensitive than amplitude to IOI, however, this is consistent with previous reports of rate effects on ABR Wave V (Hood, 1998; Lasky, 1984; Suzuki et al., 1986; Picton et al., 1981).

### 4.4 Cortical processing of frequency modulation

We evaluated cortical processing of frequency modulation via neural synchronization of EEG activity at 3.5 Hz, and sustained low-frequency activity. We previously suggested that neural synchronization, and the sustained response, although both reflecting the same stimulus, are perhaps generated in different regions of the brain and are somewhat independent (Herrmann and Johnsrude, 2018). In the present work, we found that cortical activity synchronized with the frequency modulation compared to the unmodulated control stimulus, in line with the results of previous studies (Nozaradan et al., 2011; Costa-Faidella et al., 2017; Henry and Herrmann, 2014). As for the sustained response, we observed an enhancement during the frequency modulation, compared to homologous time windows in the unmodulated control stimulus, again in line with previous literature (Barascud et al., 2016; Sohoglu and Chait, 2016; Southwell et al., 2017; Herrmann and Johnsrude, 2018; Gutschalk et al., 2002; Pantev et al., 1996).

### 4.5 Correlations of neural measures

We did not observe correlations between subcortical and cortical measures in response to our stimulus. This may imply that sensitivity to regular sounds, such as those used here, is independent at different levels of the auditory system. However, the entire auditory system is highly interconnected by dense feedforward and feedback projections (Webster, 1992; Malmierca and Hackett, 2010; Kelly and Wong, 1981; Saldana et al., 1996; Huffman and Henson, 1990) whose functions and relations are not fully understood (Oertel et al., 2002; Malmierca and Ryugo, 2011). As a consequence, responses in subcortical and cortical structures may be related indirectly or non-linearly, which makes their link more challenging to investigate using non-invasive techniques, such as the EEG-ECochG used here.

The present study did, however, observe correlations within the subcortical level. Namely, a relationship was found between the latencies of subcortical waves (Waves III and V) and their respective amplitudes. This indicates that individuals with prolonged wave latencies also display reduced wave amplitudes. This observation combined with the differential effects of IOI on the amplitudes versus latencies of subcortical responses (latency was strongly modulated by IOI, but amplitude was not) suggest that the amplitude and latency of a given subcortical response may share some, but not all, physiological underpinnings.

In contrast to the present findings, a study by Slugocki and colleagues (2017), which was able to successfully record responses from the brainstem, thalamus, and primary and secondary auditory cortices, observed cross level (subcortical to cortical) correlations between the latency of the brainstem Frequency-Following-Response (FFR) and the phase delay of the 40 Hz Auditory-Steady-State-Response (ASSR) from the auditory cortex. Additionally, they found that the amplitude of the brainstem FFR predicts the latency of the cortical N1 response. These findings suggest that subcortical and cortical processing are related. Their method, however, evaluates a different set of responses than the present study, and did not investigate adaptation of subcortical responses, or neural synchronization and sustained activity to low-frequency temporal regularity. As the relationship between levels may be complex and non-linear, it is possible that subcortical FFRs may be related to cortical ASSRs and N1, as observed by Slugocki and colleagues (2017) whereas the physiology determining the time-course of recovery from adaptation may be independent of cortical responses to regularities in sounds, such as frequency modulation. Slugocki and colleagues (2017), did not observe a relationship between subcortical responses and cortical change detection (MMN) or attention-capture (P3a) responses, consistent with the lack of correlations in the present study between subcortical and cortical responses.

### 4.6 Implications

Our novel stimulus can be used to evaluate system-wide auditory responses in humans. Our stimulus is capable of eliciting at least a subset of ABR components in addition to neural synchronization and sustained activity at cortex, allowing us to simultaneously examine sensitivity to temporal structure, and changes in temporal structure, at multiple levels. Slugocki and colleagues (2017) were able to successfully record responses from the brainstem, thalamus, and primary and secondary auditory cortices. Their findings suggest that some aspects of subcortical and cortical processing are related. They, however, evaluated a different set of responses than the present study, which study did not reveal any relationship between subcortical and cortical responses. This may be because we did not measure the appropriate characteristics or because relationships among levels are complex and nonlinear.

Our stimulus may be useful in future investigations of cognitive influences on responses in the auditory system. For example, our stimulus protocol could provide the means to investigate the effects of attention on responses throughout the auditory pathway. Investigations of effects of attention on subcortical (Rinne et al., 2008; Forte et al., 2017; Galbraith et al., 1995; Holmes et al, 2018) and cortical responses (Rif et al., 1991; Woldorff, 1993; Meltzer et al., 2015; Saupe et al., 2009; Holmes et al, 2018) have previously been conducted in isolation. This stimulus can also be applied to investigate differences in adaptation and sensitivity to changes temporal structure across groups (older vs younger adults; people with hearing impairment vs people without). Previous investigations have documented changes with age at subcortical (Willott et al., 1988; Parthasarathy et al., 2019) and cortical (Mendelson and Ricketts, 2001; Herrmann et al., 2016, 2018; Alain et al., 2014) auditory processing stages. Given these age-related changes throughout the auditory system, simultaneous measurement of responses from multiple levels might yield important information about how any coupling across levels differs in groups with different hearing abilities. While we did not observe such coupling in the current experiment, it is possible that correlations between subcortical and cortical responses would be more salient in groups with different hearing abilities, or when attention or other cognitive influences are modulated. Finally, this stimulus can be applied to investigate the relation between impaired peripheral function in people with hearing loss and subsequent maladaptive plasticity in cortical processing, such as hyper-responsivity to sound (Salvi et al., 2017; Herrmann et al., 2018; Hickox and Liberman, 2013).

## 5. Conclusion

The current study explored a novel stimulus protocol with combined electroencephalography and electrocochleography in order to assess the sensitivity of neural responses to temporal structure and changes in temporal structure at different stages of the auditory system simultaneously. Using click trains that continuously accelerate then decelerate, generating a 3.5-Hz frequency modulation (due to varying inter-click-intervals), we observed that latencies of Waves III and V, and amplitude of Wave III, of the auditory brainstem response were sensitive to inter-click-interval. At the same time, we observed neural phase-locking in the auditory cortex, and sustained activity, likely originating from higher-level cortices, to the 3.5-Hz frequency modulation. No relationship was found between subcortical and cortical responses, suggesting that any relationship between levels may be mediated by additional factors, or is potentially non-linear. By recording neural responses from different stages of the auditory system simultaneously, our research opens new avenues that enable the characterization of neural function across levels of processing, potentially enhancing sensitivity to abnormalities.

## Acknowledgements

This research was supported by BrainsCAN at Western University through the Canada First Research Excellence Fund (CFREF), by a CIHR Operating Grant to IJ, and by an NSERC Discovery Grant to IJ.

